# A highly divergent picornavirus infecting the gut epithelia of zebrafish (*Danio rerio*) in research institutions world-wide

**DOI:** 10.1101/463083

**Authors:** Eda Altan, Steven V. Kubiski, Ákos Boros, Gábor Reuter, Mohammadreza Sadeghi, Xutao Deng, Erica K. Creighton, Marcus J. Crim, Eric Delwart

**Affiliations:** Vitalant Research Institute, San Francisco, CA 94118, USA; Dept. of Laboratory Medicine, University of California, San Francisco, CA 94118, USA; Institute for Conservation Research, San Diego Zoo Global, San Diego, CA, 92112, USA; Regional Laboratory of Virology, National Reference Laboratory of Gastroenteric Viruses, ÁNTSZ Regional Institute of State Public Health Service, Pécs, Hungary; Department of Medical Microbiology and Immunology, Medical School, University of Pécs, Pécs, Hungary; Department of Virology, University of Helsinki, Helsinki, Finland.; IDEXX BioAnalytics, Columbia, MO 65201-8250, USA

## Abstract

Zebrafish have been extensively used as a model system for research in vertebrate development and pathogen-host interactions. We describe the complete genome of a novel picornavirus identified during a viral metagenomics analysis of zebrafish gut tissue. The closest relatives of this virus showed identity of ≤19.8% in their P1 capsids and ≤35.4% in their RdRp qualifying zebrafish picornavirus 1 (ZfPV1) as member of a novel genus with a proposed name of *Cyprivirus.* RT-PCR testing of zebrafish from 41 institutions from North America, Europe, and Asia showed ZfPV1 to be highly prevalent world-wide. *In situ* hybridization of whole zebrafish showed viral RNA was restricted to a subset of enterocytes and cells in the subjacent lamina propria of the intestine and the intestinal mucosa. This naturally occurring and apparently asymptomatic infection (in wild type zebrafish lineage AB) provides a natural infection system to study picornavirus-host interactions in an advanced vertebrate model organism. Whether ZfPV1 infection affects any immunological, developmental or other biological processes in wild type or mutant zebrafish lineages remains to be determined.

## Introduction

Picornaviruses are small non-enveloped viruses with positive-sense, genomic single-stranded RNA ranging in size from 6.7 kb to 10.1 kb (www.picornaviridae.com). The *Picornaviridae* family currently includes 35 genera and >75 species plus several candidate genera [1,2] and have been detected in diverse vertebrate hosts including fish (Bluegill picornavirus [3], Eel picornavirus [4], Carp picornavirus [5], Fathead minnow picornavirus [6] and unassigned fish picornaviruses [7]).

Except for the Dicipivirus genus [8,9] with two IRES and open reading frames (ORFs), typical picornavirus genomes contain a single ORF encoding a large polyprotein flanked by 5’ and 3’ untranslated regions (UTRs)[8]. The viral genome organization is as follows: the N terminal P1 region encodes the structural proteins such as viral capsid proteins and the P2 and P3 regions encode nonstructural proteins involved in polyprotein processing and genome replication.

The laboratory zebrafish, *Danio rerio,* is a member of the minnow family (*Cyprinidae*), that initially gained importance as a developmental genetics model and has since expanded into a wide range of biomedical research areas, including behavior, cancer, infection and immunology, pathogenesis, toxicology, and drug discovery. Importantly, the zebrafish has recently emerged as a valuable animal model for pathogen-host interactions for several reasons, including genetic tractability with a completely sequenced and well-annotated genome [10], a vertebrate immune system with developmental separation between innate and adaptive immune responses [11,12], the capacity to facilitate temperature-shift experiments [13], and exceptional utility for imaging, that allows infected cells and immune cells to be visualized at high resolution throughout the entire living organism [14,15]. These characteristics of the zebrafish as a host model are particularly well-suited to investigations of both viral resistance, the mechanisms that reduce pathogen burden, and the genetic determinants of viral tolerance, the mechanisms that limit pathogenesis at a given pathogen burden [14].

Several viral infection models have been established in zebrafish; however, they typically employ viruses from mammals or other fish species that did not co-evolve with zebrafish [16,17]. When infecting non-natural hosts, pathogens may display either attenuated virulence or extreme virulence [16]. Host-pathogen interactions are extremely complex, and it has been suggested that in order to preserve the intricacies of these interactions, infection models should be developed utilizing the most closely related pathogen that naturally infects the animal model species [16,18]. Investigation of virus-host interactions in a powerful animal model using a virus with a shared co-evolutionary history thus provides the opportunity to obtain a more thorough mechanistic insight into virus-host interactions.

Natural infections have also been shown to have the potential to confound experimental results in mammals [19] and zebrafish [20-22]. Research using zebrafish has the potential to be adversely impacted by unrecognized viral infections [17]. Zebrafish can also provide a model system to study vertebrate innate and adaptive responses to viruses [18] and have been infected with human viruses including herpes simplex 1 virus [23], Chickungunya virus [24], influenza A [25] and subgenomic HCV replicons [26]. To date, only two naturally-occurring viral infections have been recognized in zebrafish; Infectious spleen and kidney necrosis virus (ISKNV) in family *Iridoviridae* with linear, dsDNA genome [27], and Redspotted grouper nervous necrosis virus (RGNNV) in family *Nodaviridae* with segmented, bipartite linear, ssRNA(+) genome [28]. A lack of known naturally-occurring viral infections in zebrafish has historically been considered a disadvantage of the zebrafish as a viral host model [14]. We report the full genome sequence of a novel picornavirus with a deep root on the *Picornaviridae* family phylogenetic tree. Consistent with the nomenclature for other picornaviruses, we suggest the name genus: “*Cyprivirus”*, species: “*Cyprivirus A”*, strain: Zebrafish picornavirus-1 (ZfPV-1) for this new virus and propose that it represent the prototype of a new picornavirus genus.

## Materials and Methods

### Animals and husbandry

All zebrafish were maintained in an AAALAC-accredited facility at IDEXX BioAnalytics (Columbia, MO) in accordance with the guidelines presented in the *Guide for the Care and Use of Laboratory Animals* [29]. All husbandry and euthanasia procedures were approved by the University of Missouri—Columbia Animal Care and Use Committee. The zebrafish included 1) adult, mixed-sex wild-type AB line zebrafish sampled from a previously described breeding colony [30] originally obtained from an academic research laboratory and maintained at IDEXX BioAnalytics, and 2) adult, mixed-sex first generation offspring, exhibiting the nacre [31] phenotype, obtained from an outcross of AB zebrafish from the aforementioned breeding colony and casper [32] zebrafish that were purchased from a commercial supplier. Zebrafish were maintained on a 14:10-h light:dark cycle at 26°C in flow-through 75.7-L glass aquaria (Aqueon® Products, Franklin, WI) continuously supplied with reverse-osmosis-purified water that was remineralized with a commercially available product (Replenish™, SeaChem® Laboratories, Madison, GA), and equipped with individual air-stone aeration and biologic filtration, including power filters (AquaClear® 20 Power Filter, Rolf C Hagen, Inc., Mansfield, MA) containing plastic-foam filter material (AquaClear® 20 Foam Filter Inserts, Rolf C. Hagen, Inc., Mansfield, MA). Water quality parameters were maintained as follows: pH, 7.2-7.8; total ammonia nitrogen, 0 ppm; nitrite, 0 ppm; nitrate <40 ppm; alkalinity 40-80 ppm; and hardness, approximately 150 ppm. Zebrafish were fed at least once daily with a combination of commercial feeds (Zeigler® Adult Zebrafish Diet, Ziegler Brothers, Inc., Gardners, PA, Golden Pearls, 300-500 μm, Kens Fish, Taunton, MA).

### Euthanasia and sample collection

Zebrafish were euthanized by hypothermal shock in accordance with the AVMA Guidelines for the Euthanasia of Animals [33]. Euthanized zebrafish for viral metagenomic analysis were flash frozen in liquid nitrogen and shipped on dry ice. Euthanized zebrafish for *in situ* hybridization were flash frozen in liquid nitrogen, longitudinally bisected with a sterile scalpel, and thawed in 10% neutral buffered formalin in tissue cassettes for 24 hours prior to embedding in paraffin.

### Viral Discovery

Gut tissue from one zebrafish (line AB) was homogenized with a hand-held rotor in approximately 10X volume of PBS and frozen and thawed on dry ice 5 times. After centrifugation for 10 minutes in a table-top microfuge (15 000 x g), supernatant (400 ml) was collected and filtered through a 0.45 mm filter (Millipore). The filtrates were treated with a mixture of DNases (Turbo DNase [Ambion], Baseline-ZERO [Epicentre], benzonase [Novagen]) and RNase (Fermentas) at 37°C for 90 minutes to enrich for viral capsid-protected nucleic acids [34]. Nucleic acids were then extracted using magnetic beads of the MagMAX Viral RNA Isolation kit (Ambion) according to the manufacturer’s instructions. Random RT-PCR followed by Nextera™ XT Sample Preparation Kit (Illumina) were used to generate a library for Illumina MiSeq (2 × 250 bases) with dual barcoding as previously described [35]. An in-house analysis pipeline was used to analyze sequence data. Paired-end reads of 250 bp generated by MiSeq were debarcoded using vendor software from Illumina. Human host reads and bacterial reads are identified and removed by mapping the raw reads to human reference genome hg38 and bacterial genomes release 66 using bowtie2 in local search mode with other parameters set as default, requiring finding 60bp aligned segment with at most 2 mismatches and no gaps [36]. Adaptor and primer sequences are trimmed using the default parameters of VecScreen [37]. We developed a strategy that integrates the sequential use of various de Bruijn graph (DBG) and overlap-layout-consensus assemblers (OLC) with a novel partitioned sub-assembly approach called ENSEMBLE [38]. Sequence reads were first analyzed using BLASTx (version 2.2.7) for translated protein sequence similarity to all viral protein sequences in GenBank’s virus RefSeq database plus protein sequences taxonomically annotated as viral in GenBank’s non-redundant database using E-value cutoff of 0.01. To remove background due to sequence misclassification these initial viral hits were then compared to all protein sequences in NR using the program DIAMOND (version 0.9.6) and retained only when the top hit was to a sequence annotated as viral. To align reads and contigs to reference viral genomes from GenBank and generate complete genome sequences the Geneious R10 program was used.

The amino acid (aa) pairwise alignments and identity calculations of P1, 2C and 3CD regions were performed by the BioEdit software (ver. 7.1.3.0) using the in-built ClustalW algorithm [39,40].

The aa phylogenetic trees of picornaviruses were constructed using the Maximum likelihood method with two substitution models: Le_Gascule_2008 model (LG) with Freqs and gamma distributed, invariant sites (G + I) in case of P1 and 3CD while LG with G + I in case of 2C MEGA software ver. 7.026 [41]. The substitution models were selected based on the results of the Best Model search of MEGA 7.026. Bootstrap (BS) values (based on 1000 replicates) for each node are shown if BS > 50%.

### Genome acquisition of the new picornavirus

In order to fill two gaps in the genome of ZfPV-1 initially detected by deep sequencing, two sets of PCR primers were used for nested RT-PCR with primers designed based on the contig data. After tissue homogenization the Qiagen RNeasy mini kit was used for RNA extraction. The RT step was performed as described previously and the mixture was held at 4 °C [42].

For the initial testing for ZfPV RNA in zebrafish the primers Zf_cont1F1 (5’- ^2237^GTCGAAAACCCCAGGATCTC^2256^-3’) and Zf_cont2R1(5’- ^4221^AGTGTTGGGCTTAGAAGGGA^4202^-3’) were used for the first round of PCR, and primers Zf_cont1F2 (5’-^2266^ATGACTCTCCTCCATGCTGT^2285^-3’) and Zf_cont2R2 (5’-^4168^CTTTTCGGTAGATGAGGGCG^4149^-3’) for the second round of PCR. An initial denaturation step at 94°C for 1 minute was used followed by 40 cycles of denaturation at 94°C for 20 seconds, annealing at 54°C for 30 seconds, and elongation at 72°C for 90 seconds, with a final elongation step at 72°C for 10 minutes. Amplicons were evaluated by 2% agarose gel electrophoresis, and 1903bp product confirmed by Sanger sequencing.

The 3’ end region was completed by SMARTer® RACE 5’/3’ Kit (Takara Bio USA, Inc.) procedures were carried out according to the instructions described by the manufacturer.

The 5’ end of the virus was determined by 5’ RACE PCR (Roche Diagnostics, Mannheim, Germany) according to the manufacturer’s instructions and with minor modifications. Briefly, Maxima H minus reverse transcriptase enzyme (Thermo-Fisher Waltham, MA, USA) was used for the reverse transcription and dGTP was used for cDNA tagging [43]. The acquired single 5’ and 3’ RACE PCR products were sequenced directly using sequence-specific reverse (5’ RACE) and forward primers (3’ RACE) without the use of cloning methods. The presence of nucleotide sequence repeats was tested by Tandem Repeats Finder Program [44].

### Real-time reverse transcription PCR

To test zebrafish from other institutions submitted to IDEXX BioAnalytics, total nucleic acids were extracted from homogenized whole zebrafish using a commercially available platform (One-for-All Vet Kit, Qiagen, Valencia, California) and cDNA was synthesized using the Tetro™ cDNA Synthesis Kit (Bioline, London, UK). The ZfPV-1 real-time RT-PCR assay hydrolysis probe and primers were designed using the software PrimerExpress® version 3.0 (Applied BioAnalytics™, Waltham, MA). Real-time RT-PCR analysis was performed at IDEXX BioResearch (Columbia, MO) with standard primer and probe concentrations (Applied BioSystems™) using the master mix LightCycler® 480 Probes Master (Roche Applied Science, Indianapolis, IN) in a commercially available instrument (LightCycler® 480, Roche Applied Science). In addition to positive and negative assay controls, a hydrolysis probe-based real-time RT-PCR assay targeting a universal bacterial reference gene (16s rRNA) was amplified for all samples to confirm the presence of amplifiable DNA and absence of PCR inhibition.

### *In Situ* RNA hybridization

RNA *in situ* hybridization was performed using the RNAscope® 2.5 HD Red Chromogenic Reagent Kit according to the manufacturer’s instructions (Advanced Cell Diagnostics, Newark, CA). Target probes were designed using the custom software as described previously [45], based on nucleotides 1379-3032 of Zebrafish picornavirus 1 (ZfPV-1, GenBank accession number MH368041). Briefly, 5μm sections of formalin fixed, paraffin embedded (FFPE) tissue were mounted on AutoFrost® charged adhesion slides (Cancer Diagnostics, Inc, Durham, NC), baked at 60°C in a dry oven, and deparaffinized. The sections were treated with an endogenous peroxidase blocker for 10 minutes at room temperature before boiling in a target retrieval solution for 15 minutes. Protease plus was then applied for 30 minutes at 40°C. Target probes were hybridized for 2 hours at 40°C, followed by a series of signal amplification and washing steps. Hybridization signals were detected by chromogenic reactions using Fast Red. Slides were counterstained in 50% hematoxylin for 2 min and decolorized with 0.2% ammonium hydroxide. After rinsing in deionized water, they were dried in a 60°C oven, dipped in xylene, and cover-slipped using xylene based SHUR/Mount medium (Triangle Biomedical Sciences, Durham, NC). RNA staining signal was identified as red punctate dots. Negative control background staining was evaluated using a probe specific to a bacterial DapB gene.

## Results

Nearly one million short reads were generated from an individual zebrafish gut tissue sample processed for viral metagenomics. The raw sequence data is available at NCBI’s Short Reads Archive under GenBank accession number SRP150088. The most common detected viral reads (n=12407) in this sample belonged to the family *Picornaviridae* with BLASTx E score ranging from 7.38e^−26^ to 5.40e^−5^ to members of different genera. Following denovo assembly large contigs were generated and gaps were bridged by RT-PCR. The RNA genome extremities were acquired using Rapid Acquisition of cDNA Ends kits. Nucleic acids of another virus, avian gyrovirus 2 in the *Circoviridae* family, were also detected at the low frequency of 0.0015% (15 out of 937908 total reads).

The complete RNA genome of zebrafish picornavirus 1 strain (ZfPV-1, GenBank MH368041) is 8,298 nucleotide (nt) long excluding the poly(A)-tail (Figure 1). The genome organization is as follows: 5UTR^IRES-?/^L/P1(VP0-VP3-VP1)/P2(2A-2B-2C)/P3(3A-3B^VPg^-3C^Pro^-3D^Pol^)/3UTR-poly(A) (Figure 1). The study strain shares 49% G+C content and it has a nt distribution of 20% A, 31% U, 20% G and 29% C.

**Figure 1.**
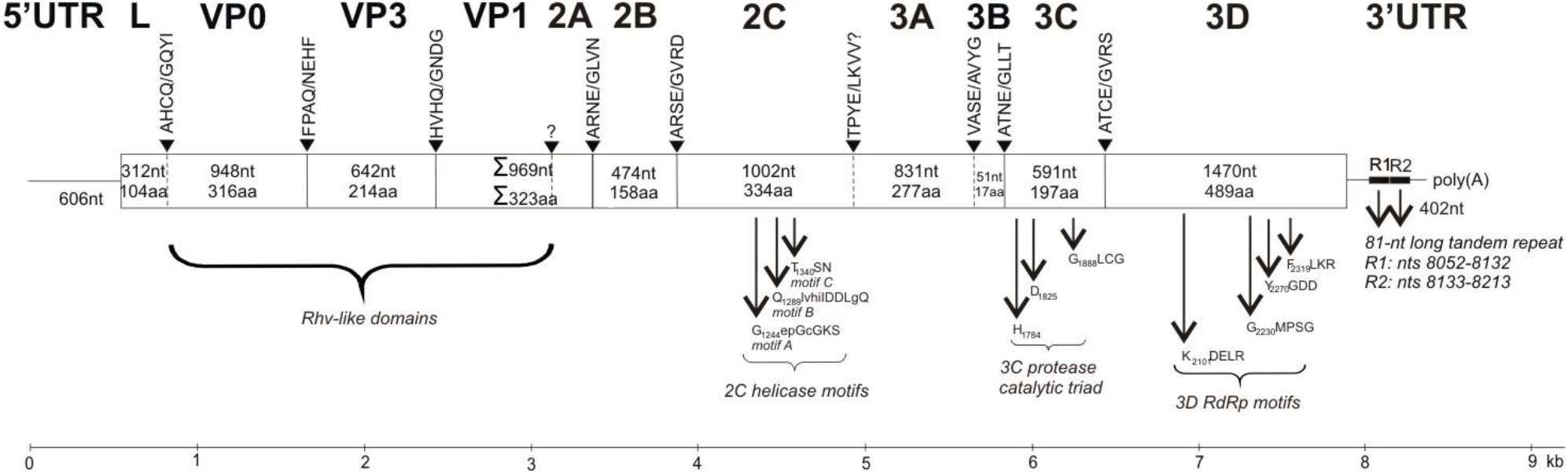
Schematic genome organization of zebrafish picornavirus 1 (ZfPV-1, MH368041). The genome map was drawn to scale. VP0, VP3 and VP1 represent viral structural proteins. Nucleotide (*upper number*) and amino acids (*lower number*) lengths are indicated in each gene box. Conserved picornaviral amino acid motifs and predicted P4/P4’ cleavage sites are indicated at the above of genome map. Question mark indicates the uncertain borders of VP1/2A and 2C/3A. The 3’UTR contains an 81-nt long tandem repeats (R1 and R2).

The 5’UTR of the study strain is 606 nt long (Figure 1). The first in-frame AUG initiation codon is at nt position 607-609 (UUUA**A_607_UG**C, start codon is bold). Similar sequences were not found in GenBank database using BLASTn and the secondary RNA structure of the 5’UTR-IRES could not be predicted: neither the five known picornavirus IRES (types I-V) structures nor the known conserved 5’UTR-IRES nucleotide motifs could be identified in the study sequence [46-48].

The predicted length of the coding region (ORF) is 7,290nt which encodes a 2,429aa long single polyprotein that is cleaved into proteins during the post-translational cleavage cascade (Figure 1). Zebrafish picornavirus 1 has a putative L-protein preceding the capsid region (Figure 1) with unknown function. The L-protein has no similar sequence in GenBank. The border of the L-protein/VP0 is potentially myristoylated (Figure 1). The predicted capsid proteins (VP0, VP3 and VP1) of zebrafish picornavirus 1 had aa identity of 25% (coverage: 82%), 34% (coverage: 99%) and 28% (coverage: 55%) to the corresponding proteins of unclassified, sapelovirus-related canine picornavirus (KU871313), Aichivirus B (NP_740257) of genus *Kobuvirus*, and feline sakobuvirus (NC_022802) of genus *Sakobuvirus*, respectively. The complete P1 of zebrafish picornavirus 1 shows the highest sequence identities (19.9%) to the corresponding genome parts of raboviruses and sakobuviruses, although only slightly lower value (19.8%) was found compared to the cardioviruses (Table 2). Consistent with the nomenclature for other picornaviruses, we propose the species name of *Cyprivirus A* and strain name of Zebrafish picornavirus 1 (ZfPV-1). Pending ICTV review this virus may also represent the prototype of a new picornavirus genus (*Cyprivirus*).

**Table 1.** Pairwise amino acid sequence identity values between the study sequence (ZfPV, MH368041) and the most closely related picornaviruses. The highest sequence identity values are in bold and underlined. Next closest identity values are in bold

| **Genus** | **Type species** | **Accession no.** | **P1** | **2C** | **3CD** |
| --- | --- | --- | --- | --- | --- |
| *Aphthovirus* | *FMDV* | NC_004004 | 16.1% | **27.8%** | 24.4% |
| *Cardiovirus* | *Cardiovirus A* | NC_001479 | **19.8%** | 22.4% | 26.1% |
| *Kobuvirus* | *Aichivirus A* | AB010145 | 18.9% | **27.8%** | 33.1% |
| *Passerivirus* | *Passerivirus A* | NC_014411 | 17.7% | 25.8% | **<u>35.4%</u>** |
| *Potamipivirus* | *Potamipivirus A* | KC843627 | 13.7% | 19.3% | 24.2% |
| *Rabovirus* | *Rabovirus A* | KP233897 | **<u>19.9%</u>** | 22.3% | 25.6% |
| *Rosavirus* | *Rosavirus A* | JF973686 | 16.5% | 25.3% | 28.1% |
| *Sakobuvirus* | *Sakobuvirus A* | KF387721 | **<u>19.9%</u>** | 25.8% | 32.7% |
| *Sicinivirus* | *Sicinivirus A* | KF741227 | 17.2% | **<u>28.0%</u>** | 33.4% |

**Table 2.**
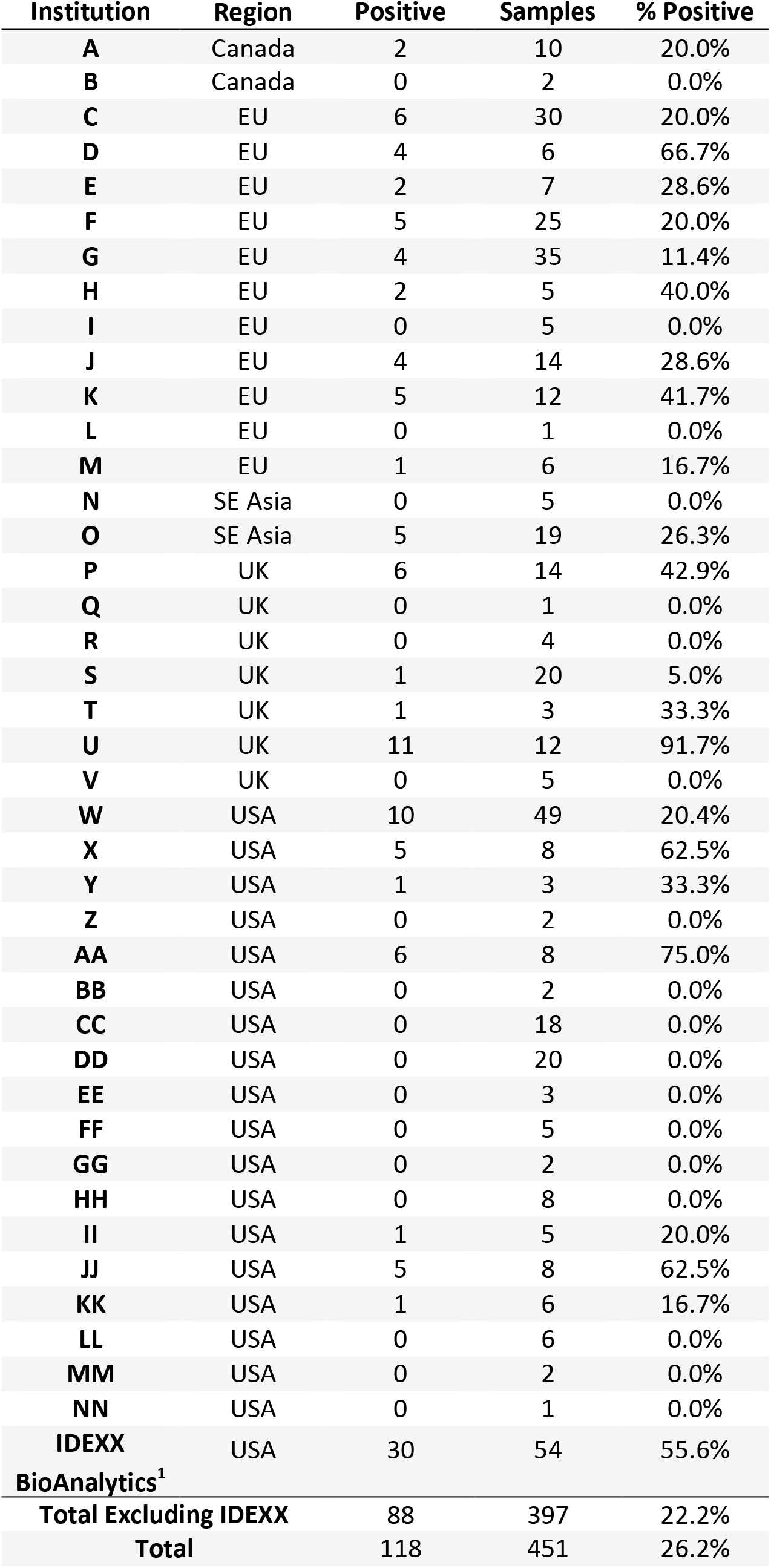
RT-PCR detection of zebrafish picornavirus and distribution across institutions. Samples represent pools of 1-5 zebrafish. ^1^All IDEXX BioAnalytics zebrafish were tested individually.

The predicted 2A is short (≈23 aa) and does not possess the characteristic catalytic aa residues of chymotrypsin-like protease, the NPGP or H-box/NC motifs. The 2C possesses the GxxGxGKS (G_1244_epGcGKS) motif for NTP binding and the D_1295_DLGQ motif for putative helicase activity (Figure 1). The picornavirus 3C protease catalytic triad and well-conserved aa motifs of 3D (RNA-dependent RNA polymerase) are present in the study strain. The 3C and 3D proteins had 33% (coverage: 78%) and 43% (coverage: 93%) aa identity to the corresponding proteins of passeriviruses (genus *Passerivirus*, NC_036588 and NC_014411), respectively. The 2C region shows the highest sequence identity (28.0%) to the corresponding genome parts of siciniviruses although only slightly lower identity values were measured (27.8%) compared to the kobu-, and aphthoviruses (Supplemental Table 1). The 3CD region shows the highest sequence identity (35.4%) to the 3CD of the passeriviruses (Supplemental Table 1).

The 3’UTR is 402 nt and similar sequences were not found in GenBank using BLASTn. Interestingly, it contains an 81-nt long tandem repeat (97.5% nt similarity to each other) from nts 8052 to 8132 and from nts 8133 to 8213 (Figure 1), respectively, with unknown secondary RNA structure and function. The 3’UTR shares high, 71% A+U content and this repeated region was confirmed twice by Sanger sequencing.

Phylogenetically, the study sequence was separated from the known picornaviruses including all fish picornaviruses in every dendrograms (Figure 2). Due to the absence of more closely related sequences the phylogenetic positions zebrafish picornavirus 1 in the P1 and 2C trees were not supported by the bootstrap analysis (bootstrap values were <50%), (Figure 2).

**Figure 2.**
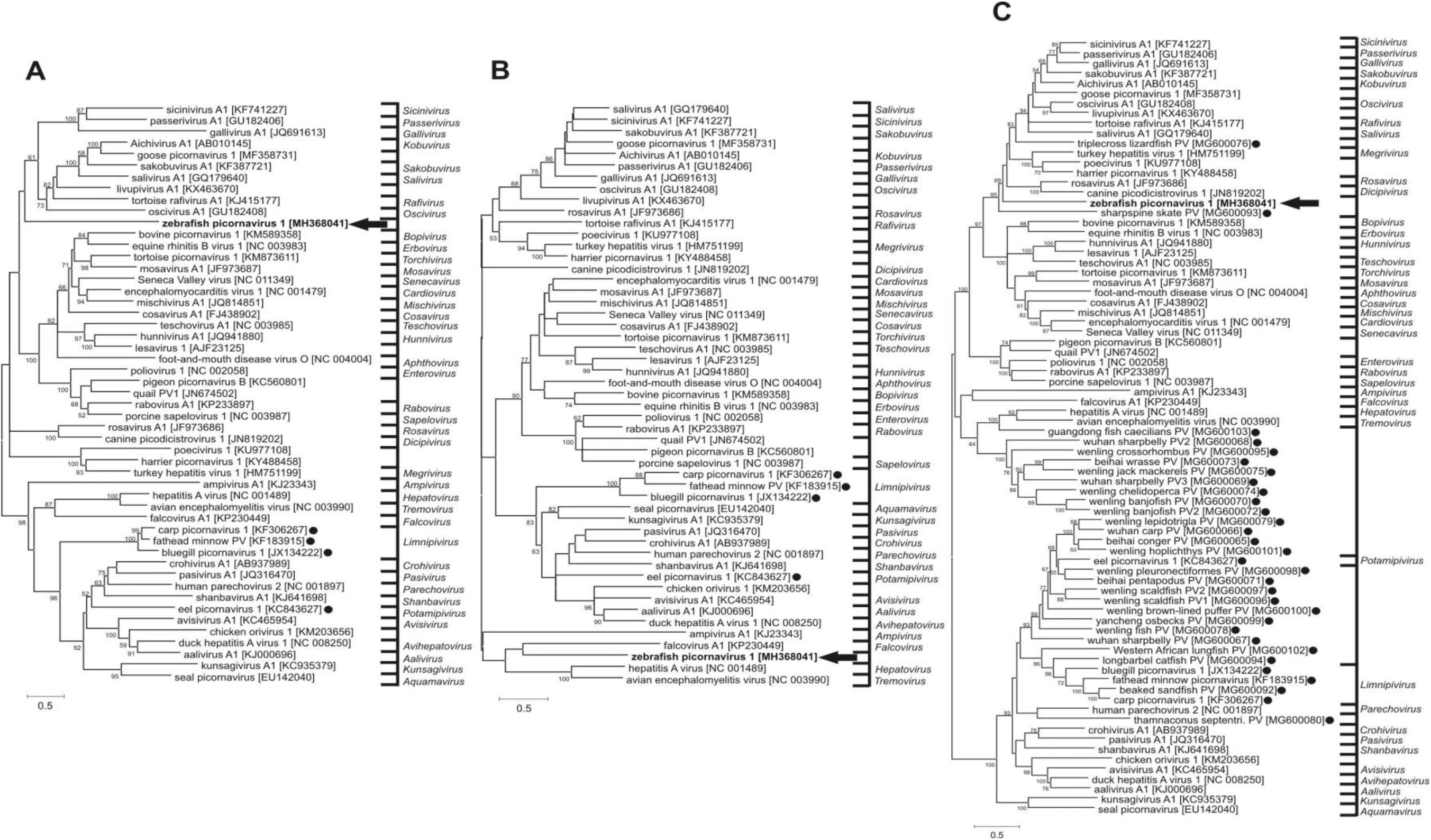
Phylogenetic trees based on the amino acid sequences of P1 (**A**), 2C (**B**) and 3CD (**C**) of zebrafish picornavirus 1 (in **bold** and marked with an arrow) and representative picornaviruses. Fish picornaviruses are highlighted with black dots. Note that the novel, currently unassigned fish picornaviruses described recently [61] were included only the 3CD tree, because the borders of P1 and 2C are not known and not adequately predict

### Detection of ZfPV RNA among IDEXX zebrafish

A nested set of PCR primers was designed to amplify a 1903 bases amplicon including a partial VP3, VP1-2A-2B-and partial 2C region. Initially 12 zebrafish without any obvious signs of disease were randomly selected from two groups of zebrafish maintained by IDEXX BioAnalytics Inc. ZfPV RNA was detected in 5 (41.6 % or 2/6 in tank 1 and 3/6 in tank 2) of 12 gut tissue samples from individual zebrafish using RT-nPCR. Positive samples were distributed as follows: 2/6 in tank 1 and 3/6 in tank 2.

### Prevalence of ZfPV in institutions world-wide

In order to estimate the prevalence of ZfPV-1 zebrafish that were submitted to IDEXX BioAnalytics for health monitoring from geographically dispersed biomedical research institutions were tested for ZfPV-1 using a real-time RT-PCR (Table 2). A total of 451 zebrafish mini-pools (1 to 5 individuals) from 41 institutions were tested. A total of 118 zebrafish mini-pools (26%) from 23 institutions (56%) were positive for ZfPV-1 RNA. Infected zebrafish were detected in all geographic regions tested (12/21 of North America, 9/11 European, 4/7 United Kingdom, 1/2 SE Asian institutions showed presence of picornavirus)(Table 2).

### Site of viral replication using in situ hybridization

Routine hematoxylin and eosin (H&E) sections showed well-preserved tissues with minimal autolysis. No pathologic lesions were observed in any of the tissues. Four fish examined with *in situ* hybridization had positive signal in the GI tract which varied in intensity between fish (Fig 3). Positive signal was interpreted as discrete, punctate red dots that were individualized to clustered or coalescent. Strong signal was consistent segmentally on the apical surface of enterocytes in all fish, within a layer of presumed mucus and digesta overlaying the mucosa, and scattered within the lumen. Signal was also seen less frequently associated with the basolateral aspect of enterocytes, within the cytoplasm, and along the basal lamina and superficial lamina propria. Both picornavirus probe and negative control probe had non-specific uptake of stain in the liver of all fish, seen as hazy, finely-granular to glassy, pink to brick-red coloration of hepatocytes. Similarly, variably-sized pieces of intraluminal debris had dense, glassy and uniform uptake of stain with viral and control probes. When present, myocardium and seminiferous tubules had similar non-specific and very faint uptake of stain. Non-specific staining was not seen in other tissues.

**Figure 3.**
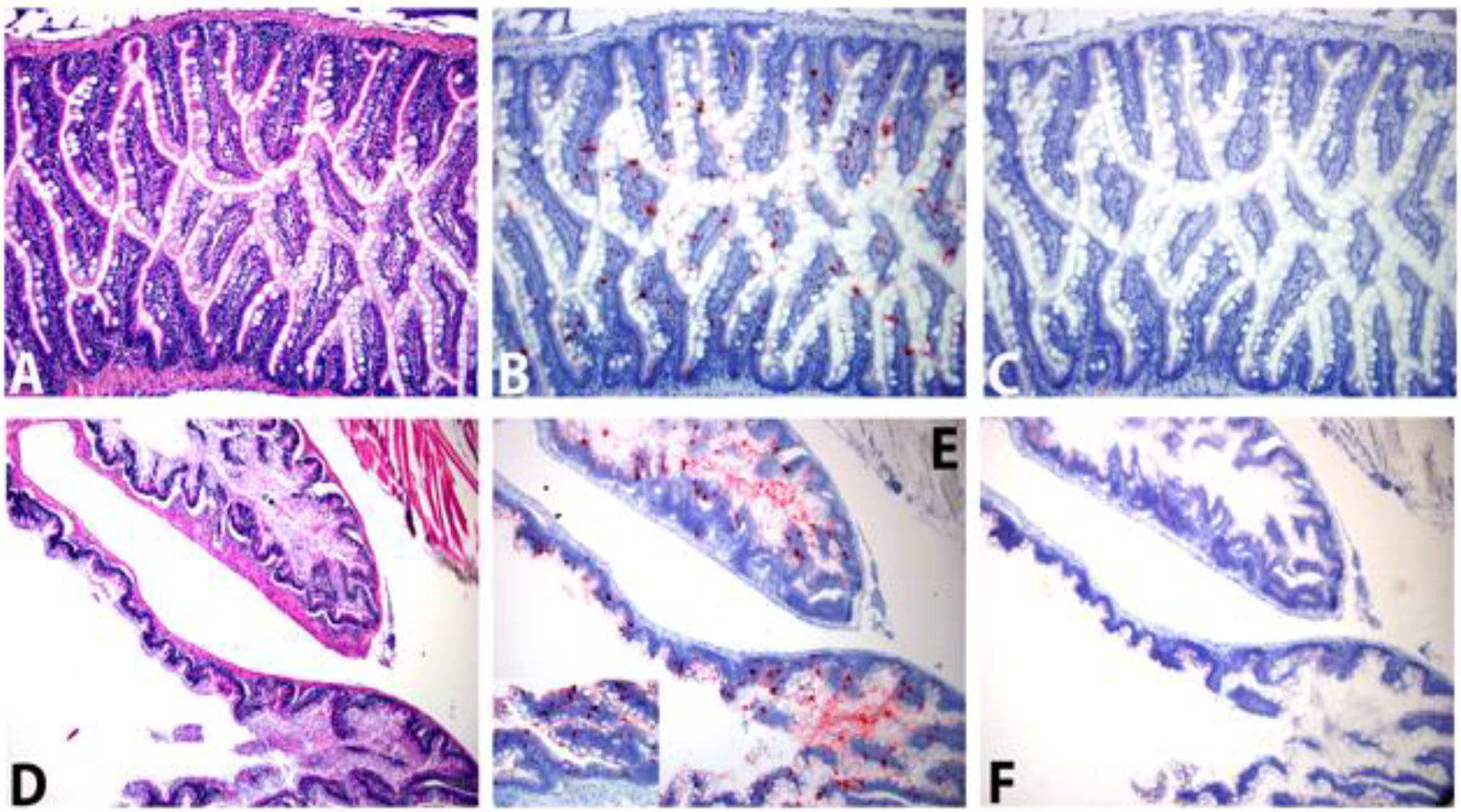
Zebrafish Picornavirus (ZfPV-1) infection. A. Longitudinal section of well-preserved intestine with minimal luminal contents and no histologic lesions. Case 5, H&E, 20X. B. Serial section at same segment as A, showing dispersed, punctate red staining of intestinal villi. ZfPV-1 probe; hematoxylin counterstain, 20X. C. Negative control for B using DapB gene probe with no positive ISH signal, hematoxylin counterstain, 20X. D. Cross section of intestinal segments. Case 1, H&E, 10X. E. Serial section at same segment as D showing similar but more densely aggregated ISH signal along mucosa and within the lumen. Hematoxylin counterstain, 10X. Inset shows higher magnification demonstrating discrete, individualized to coalescing punctate ISH signal within enterocytes, along the apical border, and scattered along the basal lamina. ZfPV probe, hematoxylin counterstain, 40X. F. Negative control for E. DapB probe, Hematoxylin counterstain, 10X

## Discussion

We describe the complete genome of a novel picornavirus (Zebrafish picornavirus 1 or ZfPV-1) detected in zebrafish. Phylogenetic trees based on the amino acid sequences of P1, 2C and 3CD showed that this genome did not cluster closely with other fish picornaviruses or any other currently sequenced picornaviruses except for the 2C region that ZfPV clustered distantly with falcovirus (KP230449) from a carnivorous bird [49,50]. Genetic distance analysis of ZfPV-1 proteins relative to those of other know picornaviruses indicates that it was sufficiently divergent based on ICTV criteria [1] to indicate the presence of a new *Picornaviridae* genus which we propose to call *Cyprinivirus* after the vertebrate family *Cyprinidae* to which zebrafish belong. Reads of avian gyrovirus 2 were also found at very low frequency. Avian gyrovirus 2 has been reported at high frequency in chicken, and may therefore originate from ingested zebrafish food products containing chicken [51-54]. The Zeigler feed used for zebrafish does include poultry by-products and hydrolyzed poultry feather meal among its ingredients.

ZfPV-1 showed a high prevalence in the initial IDEXX population in which it was initially detected. Further testing for ZfPV-1 in zebrafish from other institutions using real time RT-PCR showed that this virus was widely distributed in every geographic region tested with 21/43 institutions positive. These results likely present a minimum value as testing more samples from these institutions where ZfPV1 was not detected might have revealed a wider distribution of ZfPV1 among research institutions.

The detection of an unsuspected virus in a widely used vertebrate animal model is reminiscent of the wide spread distribution of a murine astrovirus in laboratory mice [55-57] and the presence of numerous enteric viruses in healthy rhesus macaques in primate centers [58-60]. The frequent detection of viral infections in apparently healthy experimental animals point to the need to identify and monitor such background infections when interpreting the outcomes of model animal experimentation particularly those involving innate or immune responses or developmental pathway that might be influenced by viral infections.

The absence of obvious disease signs in infected zebrafish (line AB) indicate that ZfPV-1 infection displays typically low or no pathogenicity at least in the wild type AB zebrafish lineage. Whether replication of this fish picornavirus affects zebrafish gut or immune tissue development or responses to other immunological challenges remains to be determined. Using the numerous zebrafish germ line mutations affecting different vertebrate developmental processes and innate and adaptive immune responses their role in supporting or suppressing ZfPV1 replication and possible pathogenesis can now be addressed using a naturally occurring infection. In situ hybridization showed that replication in the wild type AB strain was restricted to intestinal tissue with virus released in the luminal space providing a ready route for fecal-oral transmission of ZfPV-1, a typical mean of picornavirus transmission.

## Acknowledgements

This work was supported by Blood Systems Research Institute, and a grant from the Hungarian Scientific Research Fund (NKFIH/OTKA K111615). Á.B. is supported by the János Bolyai Research Scholarship of the Hungarian Academy of Sciences. We thank Beth A. Bauer, Cynthia L. Besch-Williford, and the necropsy and histology laboratories at IDEXX BioAnalytics, for assistance with animal husbandry and euthanasia, sample collection, and tissue processing

